# Assembly and annotation of *Solanum dulcamara* and *Solanum nigrum* plant genomes, two nightshades with different susceptibilities to *Ralstonia solanacearum*

**DOI:** 10.1101/2025.02.12.637948

**Authors:** Sara Franco Ortega, Sally James, Lesley Gilbert, Karen Hogg, Harry Stevens, Jason Daff, Ville-Petri Friman, Andrea L. Harper

**Author notes:** Corresponding author: SFO, ALH.

## Abstract

To understand why close wild plant relatives of crops, such as *Solanum dulcamara*, are resistant to *Ralstonia solanacearum* we need genetic resources to perform comparative studies and identify key genes and pathways. We *de-novo* assembled and annotated the genome of resistant *S. dulcamara* and susceptible *Solanum nigrum* plants using a hybrid approach including Oxford Nanopore Technologies and Illumina sequencing. Comparative genomic analysis was then performed to find differences between the genome of *S. dulcamara* and other susceptible Solanaceous species including potato, tomato, aubergine, and *S. nigrum* and one susceptible and one resistant *S. americanum* accession. We identified genes associated with auxin-transport only in *S. dulcamara* and a collection of pattern recognition receptors (PRRs) was identified in orthogroups only found in plant species with resistant/tolerant phenotype, suggesting novel plant receptors in these accessions that may improve recognition of pathogen-associated molecular patterns (PAMPs) associated with *R. solanacearum*. We also identified differences in methylation frequency across the gene bodies in both species, which may be associated with epigenetic regulation of resistance. Future work should assess the functional role of these PRRs during bacterial wilt development to determine if they could offer potential novel targets for breeding improved wilt resistance.

## Introduction

Bacterial wilt disease caused by *Ralstonia solanacearum* is one of the most important plant diseases worldwide, causing important economic and yield losses in more than 400 crops and being especially deadly to the Solanaceae (Mansfield et al. 2012). Despite the global efforts to find solutions to this disease, all control methods have failed to control this plant pathogen efficiently (Lemaga et al. 2001; Nicolopoulou-Stamati et al. 2016; Cardenas Gomez et al. 2024 Dec 4). Breeding for plant resistance is the most environmentally friendly and sustainable long-term solution and requires identifying quantitative trait loci (QTL) or genes associated with resistance such as the ones discovered in tobacco (Qian et al. 2013), peanut (Zhao et al. 2016), potato (Habe et al. 2019) or chilli pepper (Lee et al. 2022). However, only some resistant cultivars are available (i.e. Hawaii 7996; Wang et al. 2013), and new sources of genes associated with resistance need to be found, especially considering the additional impact climate change may have on food security and disease epidemiology. Using wild reservoir plants, such as *Solanum dulcamara*, that show low susceptibility to *R. solanacearum* is an approach that is still underexplored and could lead to the identification of new mechanisms of resistance.

*S. dulcamara* L. also called woody nightshade, climbing nightshade or bittersweet, is a semi-woody perennial belonging to the genus *Solanum* L., which is formed of more than 1400 species, including important crops such as tomato, potato or eggplant. This nightshade is predominant in the northern hemisphere, being one of the few *Solanum* sp. native to Europe, and widely extended all around the world and growing in different habitats, from riverbanks and lakes to dry areas such as dunes. *S. dulcamara* plants show high phenotypic plasticity which has caused difficulties in the taxonomy, but currently, it belongs to the Dulcamaroid clade (Huang et al. 2023), one of the 13-well supported monophyletic clades within the *Solanum* genus. Other nightshades, such as *Solanum nigrum* L. (black nightshade) are commonly found in wooded areas, gardens, vineyards and on banks (Distribution and biology of black nightshade in the UK | AHDB) and *Solanum americanum* (American black nightshade) which is considered the diploid ancestor of the hexaploid *S. nigrum* (Poczai and Hyvönen 2011), belong to the closely-related Morelloid clade. The Potato clade includes important economic crops such as potato (*S. tuberosum*) and tomato (*S. lycopersicum*) (Särkinen et al. 2013; Huang et al. 2023).

During the winter, bittersweet acts as a natural reservoir of the major plant pathogen *R. solanacearum* (Elphinstone et al. 1998; Elphinstone and Matthews-Berry 2020). Other nightshades are also important as reservoirs of other plant diseases, for example, *S. nigrum* is also known to be a reservoir of tomato yellow leaf curl geminivirus (Bedford et al. 1998).

Black nightshades (*S. nigrum* and *S. americanum*) are resistant to potato late blight (Edmonds and Chweya 1997; Lin et al. 2023) but, some *S. nigrum* accessions have recently been determined to be susceptible to *R. solanacearum* (Mafuta et al. 2022). Comparative genomic studies can help to understand the genetic basis of varying levels of susceptibility to bacterial wilt between wild and crop species. However, genomes of wild hosts are often lacking, and hence, obtaining accurate and contiguous genomes is required for comparing and identifying genes that could be linked to disease susceptibility.

In this study, we assembled the genome of a resistant/tolerant *S. dulcamara* and a susceptible *S. nigrum* accession using a hybrid approach with Oxford Nanopore Technologies and Illumina data, obtaining highly contiguous and complete genomes. We also confirmed genome size and ploidy using flow cytometry and performed de-novo annotation (including transposable elements) of the new genomes using mRNA data from different tissues. The annotations allowed comparative genomics analysis based on gene orthologues, confirming the validity of the annotation of these two new genomes and identifying genes present only in those plant species with low susceptibility to *R. solanacearum*. In *S. dulcamara*, there were genes with functions related to auxin-transport. We also identified pattern recognition receptors (PRRs), involved in recognition of pathogen-associated molecular patterns (PAMPs) in two resistant species, suggesting an arsenal of genes that could facilitate pathogen recognition and activation of the plant immune system. Finally, as Oxford Nanopore Technologies also allows DNA methylation to be quantified, we assessed how methylation changes across different genomic locales in both genomes and in genes identified only in less susceptible plant species. We also associated methylation frequency patterns with transposable elements.

## Material and methods

### Plant growth

*S. dulcamara* and *S. nigrum* seeds were obtained from the Millenium Seed Bank, Royal Botanic Gardens Kew (ID-39084 isolated from the UK in 1982 and ID-170516, isolated from the UK in 2000). The plants were grown and propagated at the University of York (York, UK). Seeds were surface sterilised for 1 minute in bleach/water (1:99 v/v), rinsed with distilled water and then stratified for 4 days on wet tissue paper at room temperature. After germination, the seeds were moved to compost (John Innes No2, approximate 125 g compost/pot) in a growth room kept at 20°C (±2°C) and with 14h/10h light /dark conditions with the lights (Model L28, Valoya, Loughborough, UK) suspended approximately 40cm above the crop which provide a light intensity of about 120 µmol m^−2^s^−1^ (SD ±10). The plants were self-pollinated to obtain berries.

### Plant inoculations with *R. solanacearum*

To assess the susceptibility of both *S. dulcamara* and *S. nigrum* to *R. solanacearum*, plant inoculations were performed and compared to a susceptible tomato cultivar “Moneymaker” (Franco Ortega et al., 2024). The three plant species were grown as above for 17 days. At 17 days (4 days before the inoculation), plants were moved to a PHCbi growth cabinet (24°C 16 h light, 20°C 8h dark, with light intensity reaching 220 µmol m^−2^s^−1^ measured from the centre of the cabinet (3 Panasonic FL40SSENW37 fluorescent tubes (37W/5700K)/side on 3 sides of the cabinet) to acclimate. Bacterial inoculum was prepared by growing *R. solanacearum* Race 3 Biovar 2 strain UW551 stock in CPG liquid media (Casamino acid 1g/L, Peptone 10g/L, Glucose 15g/L) at 28°C for 3 days, maintaining agitation at 100 rpm. 24 hours before plant infections, fresh CPG sterile media (10% of the volume of the inoculum) was added to the bacterial cultures. The OD_600_ was adjusted to 0.7 (8.3E+09) before 5 mL of the bacterial inoculum was added as a root drench (21 day-old plants; N=18/plant species). The same number of plants were maintained as negative controls by adding 5 mL of sterile CPG media onto roots. The symptoms were recorded for 21 days post-infection using a scale of 0-4, where 0=plants with no wilting symptoms, 1=25% of the plant surface showing symptoms, 2=50% of the plant showing wilting symptoms, 3=wilting symptoms in 75% of the plant and 4=dead plants. Curves of bacterial wilt disease progression and bar plots reporting the number of healthy (DI=0) or diseased plants (DI>0) for each plant species were plotted using ggplot2 package in R.

### Plant material for genome assembly

DNA was extracted from 3-month-old adult plant leaves using the high molecular weight gDNA extraction using the protocol described in Vaillancourt and Buell (2019) which uses Carlson buffer and the QIAGEN Genomic-tips 500/G (QIAGEN, Manchester, UK). The gDNA quality was assessed with a Nanodrop and Agilent Tapestation 4200 (Agilent, Cheadle, UK). The *S. dulcamara* gDNA library was prepared using Oxford Nanopore Technologies’ (ONT) ligation library preparation kit LSK109 and run using R9.4.1 flowcells on a PromethION 24 by the Bioscience Technology Facility, York. The same gDNA was also sequenced using the NovaSeq 6000 platform (Illumina, CA, USA) with a paired-end library strategy (PE150) provided by Novogene UK (Cambridge, UK). The *S. nigrum* genome was obtained using a ONT’s ligation kit LSK114 and run on PromethION R10.4.1 flowcells, keeping only the high accuracy reads following superaccuracy basecalling with guppy software version 6.4.6.

For *de-novo* annotation, we collected roots, stems, leaves, flowers and berries from each plant at the same time as the DNA samples, and these were stored at –70°C prior to RNA extraction. The plant material was ground in liquid nitrogen and the RNA extracted using the E.Z.N.A. ® Plant RNA Kit (VWR, Lutterworth, UK), which included DNase treatment to remove residual DNA according to the manufacturer’s instruction. The quality of the RNA was assessed with Nanodrop, Qubit 4.0. (Thermo Fisher Scientific, Milton Park, UK) using the Qubit™ RNA Broad Range kit (Thermo Fisher Scientific) and the Agilent Technology 2100 Bioanalyzer (Agilent Technologies, CA, USA). Only samples with RNA integrity number >6 were used to prepare the cDNA libraries. For the *S. dulcamara* genome annotation, full length barcoded cDNA libraries were generated for each tissue using ONT’s cDNA barcoding kit PCB109, and barcoded cDNA libraries were pooled for running on a R9.4.1 flowcell on a PromethION 24. The same RNA was used to for short read sequencing by Novogene services (Cambridge, UK) using the NovaSeq 6000 platform, a paired-end strategy (PE150) and an Illumina mRNA library preparation (poly A enrichment). For the *S. nigrum* genome annotation, we obtained RNA from the same tissues collected at the same time points as with *S. dulcamara* but sequenced them only with the Illumina NovaSeq 6000 platform, using equimolar concentrations of the 5 pooled tissues.

### Chloroplast and mitochondrial genome assembly

The *S. dulcamara* chloroplast assembly was started by using minimap2 to map the raw ONT reads against the complete *Solanum lycopersicum* chloroplast (NC_007898.3), followed by *de-novo* assembling the mapped reads using CANU (version 2.1.1) (Koren et al. 2017) with setting genomeSize=155k. RACON version 1.4.20 and MEDAKA version 1.0.3 were further used to polish the first preliminary chloroplast assembly (Vaser et al. 2017). To improve the assembly, the same ONT reads were mapped again against this preliminary chloroplast assembly followed by a second round of assembly using CANU, RACON and MEDAKA. Unmapped reads obtained after the second mapping round for the chloroplast, were then mapped using minimap2 against the *Solanum lycopersicum* mitochondrion complete genome (NC_035963.1), and mapped reads were used to assemble the first preliminary *S. dulcamara* mitochondrial genome following the same approach with CANU, RACON and MEDAKA. As with the chloroplast assembly, this preliminary mitochondrion assembly was followed a second round of assembly as before.

The *S. nigrum* chloroplast and mitochondria were directly assembled using minimap2 by mapping the raw ONT reads to the *Solanum lycopersicum* chloroplast (NC_007898.3) and *Solanum lycopersicum* mitochondrion complete genome (NC_035963.1), with one round and two rounds of assembly with CANU, RACON and MEDAKA, respectively.

### Flow cytometry for determining genome size and ploidy with *S. dulcamara* and *S. nigrum*

To assess the nuclear genome size and ploidy of both *S. dulcamara* and *S. nigrum*, we used flow cytometry following the protocol by Doležel et al (2007) using TrisMgCl_2_ buffer for the homogenization step. *Solanum lycopersicum* (tomato), and *Zea mays* (maize) were used as references to estimate the genome sizes. The protocol was performed with a total of 20 mg of leaves using 10 mg of the target and 10 mg of the reference.

### Nuclear genome assemblies

Different approaches were used to assemble the nuclear genomes for each plant species to obtain the most contiguous and complete genome assemblies. After assembly of the chloroplast and mitochondrial genomes, CANU (version 2.1.1; Koren et al., 2017) was used to assemble the nuclear genome using the unmapped reads. To phase the genome, PURGE HAPLOTIGS (Roach et al. 2018) reassigned contigs to haplotigs according to read depth and homology. Finally, the *S. dulcamara* genome was polished with Illumina data using PILON version 1.24 (Walker et al. 2014). The *S. nigrum* genome was assembled with FLYE (Kolmogorov et al. 2019) using the high-accuracy reads from the PromethION R10.4. run, followed by RACON version 1.4.20 and MEDAKA version 1.0.3. (Vaser et al. 2017) to polish the genome using the same high accuracy reads. Lastly, PURGE HAPLOTIGS was used to reassign contigs to haplotigs.

SEQKIT was used to assess the assembly size and N50. Completeness of the genome was assessed with the Benchmarking Universal Single-Copy Orthologs (BUSCO version 5.5; Manni et al., 2021) against the Solanales and Viridiplantae databases. ^1718^

To annotate the genomes, repetitive elements were identified using REPEATMODELER (version 2.1) and REPEATMASKER (version 2.1; Flynn et al., 2020). BRAKER (version 1.9) (Brůna et al. 2021) was used to predict the gene models in the masked genome. Protein data from the Viridiplantae (downloaded November 2021; https://v100.orthodb.org/download/odb10_plants_fasta.tar.gz; Zdobnov et al., 2021) was used in combination with ONT and Illumina mRNA-Seq data for *S. dulcamara,* and only Illumina for *S. nigrum*. TSEBRA (Gabriel et al. 2021) with keep_ab_initio configuration was used to combine the results of both proteins and mRNA-Seq predictions. Annotations were filtered by structure and function using GFACS (Version 1.1.2); Caballero and Wegrzyn, 2019) and ENTAP (Hart et al. 2020). EggNOG (Huerta-Cepas et al., 2019; Cantalapiedra et al., 2021) was used for the functional characterization of the gene models.

### Identification of methylation patterns and transposable elements in plant genomes

To identify potential epigenetic regulation in these wild species, we assessed DNA methylation and methylation frequency of the *S. dulcamara* and *S. nigrum* genomes using f5c v1.2 (Gamaarachchi et al. 2020) setting the “pore” option to r9 or r10. The frequency was only calculated when the log_lik_ratio (log-like methylated – log-like unmethylated) had a positive value, supporting methylation. Transposable Elements (TE) and tandem repeats were identified using the Extensive de-novo TE Annotator (EDTA; Ou et al., 2019). For each plant species, the methylation frequency was assessed across three genomics locales: the gene body, 1 kbp upstream and 1 kbp downstream considering all the genes in the genomes. TEs were classified into terminal inverted repeat sequences (TIRs), non-TIRs or long terminal repeats (LTRs). Plots representing average methylation frequency across the genomic locales in function of the TE type (or genomic locales without TEs), were obtained by dividing the gene body length into 20 intervals each representing 5% of the gene’s length, whilst the upstream and downstream regions were divided into 20 intervals corresponding to 50bp each. Methylation frequency was also assessed for specific genes identified during the comparative genomics analysis. To assess if the methylation frequency was conserved between these two plant species, we performed Pearson correlations between the methylation frequency of *S. dulcamara* and *S. nigrum* in the three genomic locales and plotted them with ggscatter from the R package ggpubr (Kassambara A 2023).

### Comparative genomics to find orthogroups and shared genes between plant species with different susceptibility to *R. solanacearum*

To find genes present only in plant species with lower susceptibility to bacterial wilt, we used the nightshade genomes assembled here, along with the genomes of *S. tuberosum* C88 (http://spuddb.uga.edu/data/), tomato (pangenome of *S. lycopersicum;* Zhou et al., 2022), aubergine (*S. melongena* (https://solgenomics.net/ftp/genomes/Solanum_melongena_V4.1), and two accessions of *S. Americanum:* SP2773 and SP2775 (Moon et al. 2021). The 7 genomes were inputted into the Synteny Imaging tool (SYNIMA; Farrer, 2017) using the Orthofinder pipeline (Emms and Kelly 2019). The unrooted species tree was produced using the STAG (Species Tree inference from All Genes) algorithm (Emms et al. 2018 Feb 19) using the command Orthologs_to_summary.pl. After obtaining the Orthogroups.csv file, we used kinfin (version 3; Laetsch and Blaxter, 2017) to calculate the number of orthogroups shared between the plant species. We also retrieved genes belonging to orthogroups only present in *S. dulcamara,* and those also shared with the resistant *S. americanum* SP2773 accession. To understand the function of these genes only present in species with low susceptibility to bacterial wilt, the *S. dulcamara* genes in these orthogroups were retrieved and a gene ontology (GO) enrichment analysis was preformed using the topGO package (Bioconductor – topGO) in R, using the whole genome annotation of *S. dulcamara* as the universe. The R package UpSetR (Conway et al., 2017) was used to plot the orthogroups shared between species.

## Gene expression of selected pattern recognition receptors

The mRNA-Seq used for the *S. dulcamara* annotation was also used to evaluate the expression of selected pattern recognition receptors (PRRs) (Sd_g3085.1, Sd_g3087.1, Sd_g3090, Sd_g26139.3, Sd_g8870, Sd_g28574), identified during the comparative genomic analysis with other plant species. Salmon v.1.10 (Patro et al. 2017), was used to obtain transcript-per-million (TPMs) in each of the 5 plant tissues (roots, stems, leaves, flowers and berries) using the long ONT reads and default parameters. The expression of these genes was also compared with other PRRs (leucine-rich repeat receptor-like kinases; LRRs). Plots of log (TPMs+1) were obtained using ggplot2 in R.

## Results and discussion

### Comparing plant susceptibility to *R. solanacearum*

We first compared the susceptibility of *S. dulcamara*, *S. nigrum* and tomato to *R. solanacearum* infections. Only 1 out of 18 *S. dulcamara* plants showed bacterial wilt symptoms at 21 dpi, compared with 9 plants showing wilting symptoms in tomato cv Moneymaker, 3 of them with disease index 4 and the rest with disease index 3 at 21 dpi (Fig. 1A and Fig. 1B). In contrast, at 21 dpi, *S. nigrum* showed high susceptibility to *R. solanacearum* with 15 out of 18 plants showing bacterial wilt symptoms, including one completely dead plant (DI=4) and 7 plants with a disease index of 3 (Fig. 1B). Overall, this experiment confirmed that *S. nigrum* and *S. dulcamara* have different levels of susceptibility to *R. solanacearum* at 24°C. *S. dulcamara* showed almost no symptoms after three weeks post-infection confirming tolerance or partial resistance, as reported previously by Sebastià et al. (2021).

**Fig. 1.**
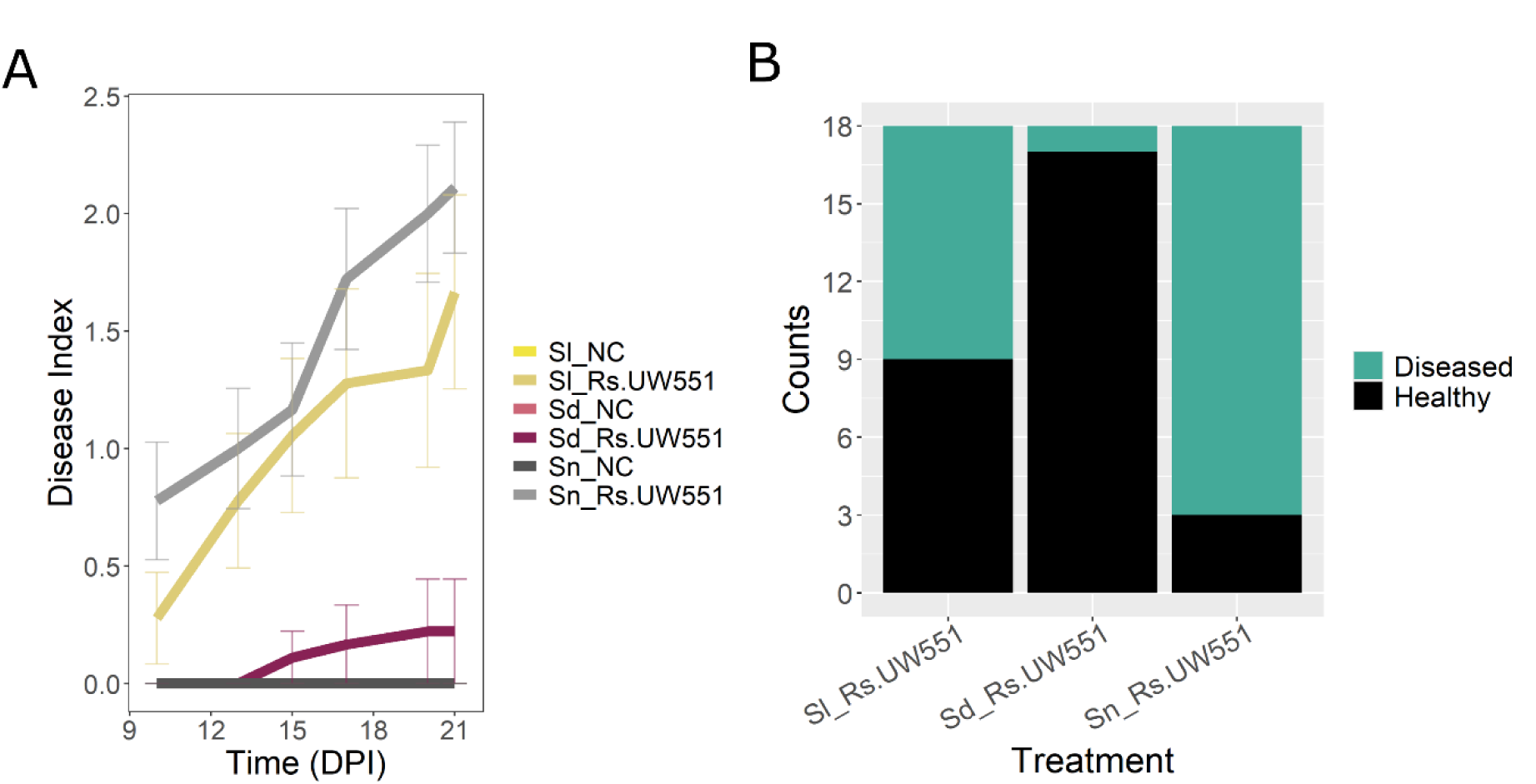
Disease index at 21 days post-inoculation of *S. dulcamara, S.nigrum* and tomato plants confirming the resistance/tolerance mechanisms of nightshades against *R. solanacearum* at 24°C. **A**. Disease index at 21 dpi. **B**. Number of diseased and healthy plants in each treatment. **C.** Area under the disease curve.

### Genome assembly and annotation

To identify genes or pathways associated with different susceptibility to *R. solanacearum*, we started by de-novo assembling the genome of the partially resistant *S. dulcamara* and susceptible *S. nigrum*. We started by assessing genome size with flow cytometry, followed by hybrid assembly of the genome using ONT and Illumina data.

Flow cytometry provided similar estimates of diploid genome size for the two nightshades: 1.03 Gbp for *S. dulcamara* and 1.29 Gbp for *S. nigrum.* Recently Christenhusz (2023) reported a *S. dulcamara* genome size of 1.2Gbp by flow cytometry.

The *S. dulcamara* chloroplast genome was assembled into a single contig with 226,924 bases (>200x coverage considering a size of 155 kbp) whilst the mitochondrial genome was assembled using 2,785 reads (91,027,879 bases; >200 times coverage) into 18 contigs covering a total length of 457,235 bp (N50=54,728 bp).

After mitochondrial genome assembly, the remaining 1,048,676 reads (26,787,058,484 bases, N50=34,429) provided 26x coverage of the nuclear genome, based on the genome size estimated using flow cytometry. After phasing and polishing with Illumina data, the genome assembly constituted a total of 3,356 contigs covering 881,223,639 bp (N50=421,481bp). BUSCO scores reached 89.9% for Solanales and 94.4% for Viridiplantae databases (Table 1).

**Table 1.** BUSCO scores of the new *S. dulcamara* and *S. nigrum* genomes.

|  | *S. dulcamara* |  |  |  | *S. nigrum* |  |  |  |
| --- | --- | --- | --- | --- | --- | --- | --- | --- |
|  | Viridiplantae |  | Solales |  | Viridiplantae |  | Solales |  |
|  | Numb<br>er | Percenta<br>ge | Numb<br>er | Percenta<br>ge | Numb<br>er | Percenta<br>ge | Numb<br>er | Percenta<br>ge |
| **Complete** | 401 | 94.40% | 5349 | 89.90% | 423 | 99.60% | 5691 | 95.70% |
| **BUSCOs**<br>**(C)** |  |  |  |  |  |  |  |  |
| **Complete**<br>**and**<br>**single-**<br>**copy**<br>**BUSCOs**<br>**(S)** | 391 | 92.00% | 5394 | 86.40% | 407 | 95.80% | 5513 | 92.70% |
| **Complete**<br>**and**<br>**duplicate**<br>**BUSCOs**<br>**(D)** | 10 | 2.40% | 208 | 3.50% | 16 | 3.80% | 178 | 3.00% |
| **Fragment**<br>**ed**<br>**BUSCOs**<br>**(F)** | 8 | 1.90% | 65 | 1.10% | 1 | 0.20% | 55 | 0.90% |
| **Missing**<br>**BUSCOs**<br>**(M)** | 16 | 3.70% | 536 | 9.00% | 1 | 0.20% | 204 | 3.40% |
| **Total**<br>**BUSCO**<br>**groups**<br>**searched** | 425 |  | 5950 |  | 425 |  | 5950 |  |

The *S. nigrum* genome was assembled using 83,200,377,815 bp (N50=19,765, 69x coverage), obtaining 3,010 contigs, totalling 1,061,442,536 bp (N50=1,883,648). BUSCO scores reached 95.7% for Solanales and 99.6% for Viridiplantae databases (Table 1). The *S. nigrum* chloroplast genome was assembled into 7 contigs (N50= 190,205) with 2094 reads (31,004,936 bases; ∼200x coverage) consisting of 263,207 bases. The mitochondrial genome was assembled into 4 contigs (N50=102,821) using 5081 reads (90,008,286 bases; ∼200x coverage considering a size of 450 kbp) and resulting in a final length of 396,930 bp.

The GC content average was 35.65% for *S. dulcamara* and 36.54% for *S. nigrum.* Despite the *S. dulcamara* genome reported here being assembled into a higher number of contigs than the genome recently reported by Christenhusz (2023), with a total length of 946.3 Mb in 105 sequence scaffolds, we additionally performed de-novo annotation obtaining a total of 27,429 genes (23,800 with functional annotation), which expanded on the *S. dulcamara de novo* transcriptome of 24,193 unigenes reported by (D’Agostino et al. 2013). On the other hand, the number of annotated genes for *S. nigrum* reached 39,466 (30,426 with functional annotation), which was fewer than the transcriptome of 47,470 unigenes reported by (Heo et al. 2022) using a *S. nigrum* obtained from NIBR, Incheon, Republic of Korea. This difference might indicate high diversity between isolates/cultivars from different geographical origins and indicates the necessity of using pangenome approaches to obtain comprehensive gene repertoires in wild species.

### *S. nigrum* showed higher DNA methylation frequency across the genome

Epigenetic changes such as DNA methylation are regulatory mechanisms that play a key role in plant-pathogen interactions, for example by modulating gene expression of defense-related genes (Wu and Fan 2025 Jan 25). We started by assessing whole genome methylation frequencies in both plants and associating these frequencies with the type and content of transposable elements (TEs) as they can be associated with silencing (due to hypermethylation) of the genes they are located inside or next to (Lee et al. 2023). The methylation frequency in the *S. dulcamara* contigs was 77.4% with an average of 23.4 called sites per contig and 18.11 methylated called sites (Table S1). The methylation average increased to 87.5% methylation frequency on the *S. nigrum* genome (with an average of 15.9 methylated sites out of an average of 34.9 called sites; Table S2). When the methylation frequency was assessed across different gene locales, it followed similar patterns for both plant species, but it was 17.1% higher in *S. nigrum* than in *S. dulcamara* in the 1 kbp upstream region (67.3% vs 50.2% average), 13.7% in the gene body (78.9% vs 65.9%), and 21.5% 1 kbp downstream region (71.3% vs 49.9%) (Fig. 2A). Using different ONT flow cells might explain these differences, however, Ni et al. (2023) recently confirmed the average proportion of 5mC called using R9.4.1 and R.10.4 flow cells is similar (67.44% and 67.70%, respectively), with strong Pearson correlation (0.64) between the results of both in various gene locales. However, the authors confirmed that R10.4 showed a lower background than R.9.4.1 in regions with high GC content Ni et al. (2023).

**Fig. 2.**
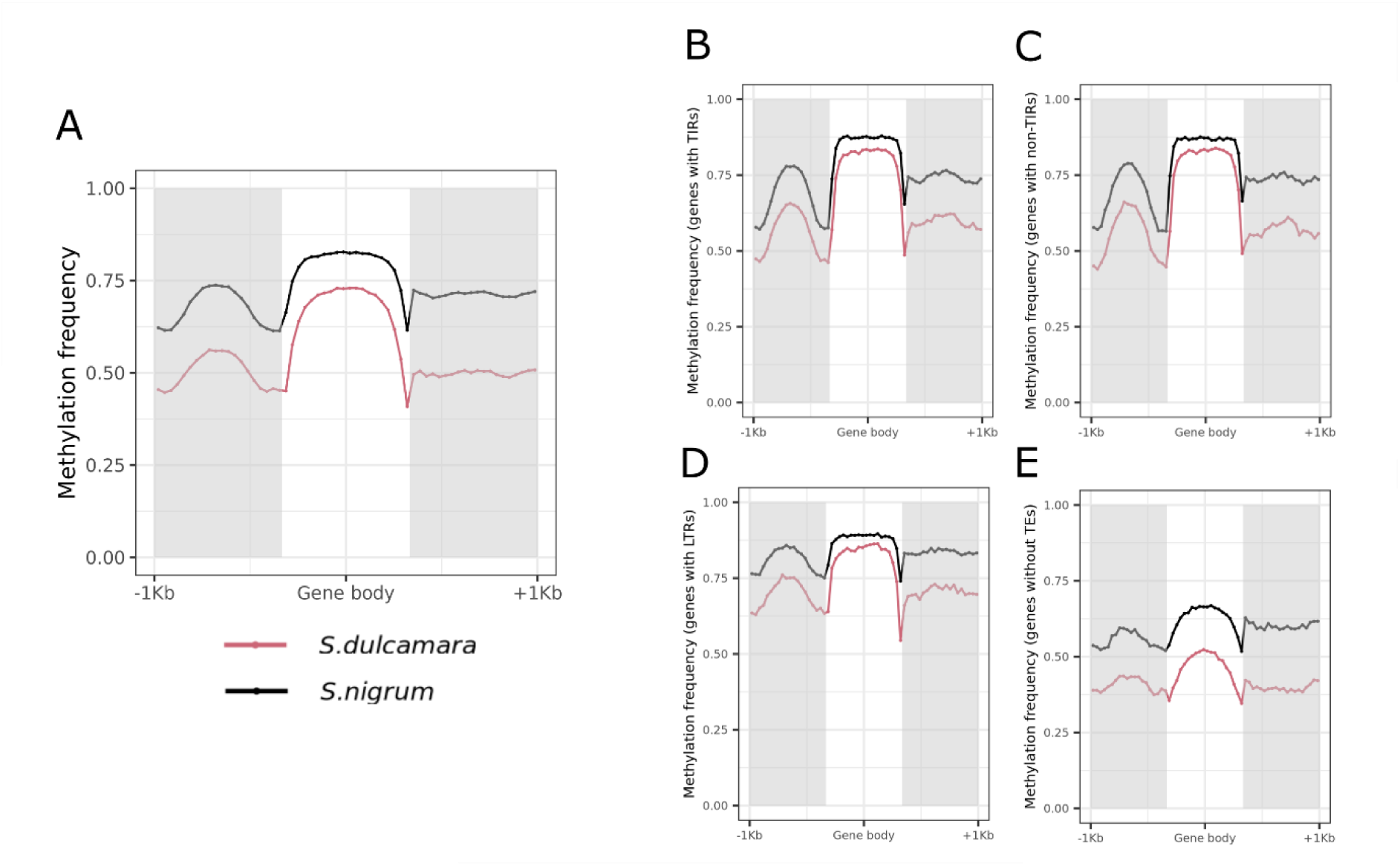
Methylation and TEs across gene body and upstream and downstream regions considering all the genes in the genomes. **A**. Methylation frequency across gene body, 1 kbpp upstream and 1 kbpp downstream for all the *S. dulcamara* (pink) and *S. nigrum* (black) genes. **B – D.** Methylation frequency across gene body, 1 kbpp upstream and 1 kbpp downstream for all the *S. dulcamara* (pink) and *S. nigrum* (black) genes containing TIRs (B), non-TIRs (C) and LTRs (D) and genes without TEs (E).

Considering the Ni et al. (2023) results, we concluded the differences in methylation frequency between both genomes can only be partially explained by using different flow cells for sequencing. Instead, the overall higher methylation frequency in the *S. nigrum* genome could be associated with different epigenetic regulation, for example, modulation by transposable elements (TEs), which are usually repressed by epigenetic silencing via hypermethylation. TEs also constitute a large part of plant genomes and are essential for the adaptative evolution of a species (Wyler et al. 2020; Chu et al. 2024).

We confirmed that the percentage of TEs was lower in *S. dulcamara* (948,592 TEs masking 62.86% of the genome) than in *S. nigrum* (881,028 TEs masking 72.94% of the genome). When looking at the distribution of TEs in different genomic locales (gene body, 1 kbp upstream and 1 kbp downstream areas considering all genes in the genome), we observed that *S. nigrum* had more TEs than *S. dulcamara* in all locales. On average across the 27,429 genes in *S. dulcamara*, we observed 1.18±1.24, 2.19±4.26 and 0.96±1.16 TEs/gene for upstream, gene body and downstream regions, respectively. These numbers increased to 1.67±1.50, 2.87±5.12 and 1.4±1.41 TEs/gene for upstream, gene body and downstream regions, respectively in the *S. nigrum* genome (39,446 genes). As TEs are often hypermethylated, greater numbers of TEs in the *S. nigrum* genome may explain the higher methylation frequency observed in comparison to *S. dulcamara*.

However, higher methylation frequencies in *S. nigrum* for all TE types in gene bodies were also observed (78.2% in *S. dulcamara* vs 85.1% in *S. nigrum* for gene bodies with TIRs; 78.4% vs 85% for gene bodies with non-TIRs and 80.9% vs 87.3% for gene bodies with LTRs; Fig. 2B, 2C and 2D), suggesting that not only are there more TEs in the *S. nigrum* genome, but they are generally more highly methylated in this species. Similar patterns were observed in upstream regions (55.7% vs 67.6 % for upstream regions with TIRs; 54.8% vs 67.5% for upstream regions with non-TIRs and 69.2% vs 80.6% for upstream regions with LTRs), and downstream regions (59.5% vs 74.2 % for downstream regions with TIRs; 56.9% vs 73.8 downstream regions with non-TIRs and 69.2% vs 83.5% for downstream regions with LTRs; Fig. 2B, 2C and 2D). In genes without TEs, methylation frequency was reduced to 46.1% and 62.6% in gene bodies, 40.7% and 55.4% in the upstream region and 39.9% and 60.2% in the downstream region for *S. dulcamara* and *S. nigrum*, respectively. This indicates that the increase in methylation observed in *S. nigrum* is a general feature which is not restricted to TEs.

The methylation frequency of all genes was positively correlated in these two plants, meaning they followed similar methylation frequency patterns despite the higher percentage methylation observed in the *S. nigrum* genome. All Pearson correlations were significant, but they were stronger for the upstream and gene bodies locales for the three TE types than the downstream regions or in genes without TEs (Fig.S1), suggesting both upstream and gene bodies showed conserved methylation patterns. These results confirmed that the higher methylation observed in *S. nigrum* (Figure 2A) was likely not due to different ONT flow cells used for sequencing but could be associated with both a greater accumulation of TEs in *S. nigrum*, and increased methylation of TEs within gene regions.

Overall, we obtained two new contiguous genomes and showed differences in methylation frequencies across the genomic locales that might be linked with higher TE percentage in *S. nigrum* than in *S. dulcamara*. Differences in epigenetics might have adaptative and evolutionary consequences that should be explored in future work.

### Comparative genomics confirms the presence of genes related to immunity and phytohormones only in resistant/tolerant accessions

To identify genes associated with bacterial wilt resistance/tolerance, we performed a comparative genomics analysis with 7 plant species (resistant/tolerant species: *S. dulcamara* and *S. americanum* SP2773 and susceptible species: *S. nigrum*, *S. americanum* SP2775, *S. lycopersicum*, *S. melongena* and *S. tuberosum*) focusing on the Orthofinder results. A total of 449,192 genes were evaluated, assigning 398,266 genes to 29,498 orthogroups and 50,926 genes not being assigned to orthogroups.

A high percentage of the genes (>83%) (Table S3, Table S4, Table S5) in each species were assigned to an orthogroup, whilst unassigned genes, those that did not produce a good BLAST hit against the full transcriptome dataset using the default SYNIMA Blast_grid_all_vs_all.pl script parameters were relatively few, indicating that most genes had been retained in at least some of the species. In total, we obtained 12,574 (42.6%) orthogroups with all species present, 5,549 species-specific orthogroups (which contained multiple paralogs in one species), and 51 single-copy orthogroups (ie. containing exactly one gene copy in each species) (Table S4). The median size was 10 genes per orthogroup.

A high number of orthogroups (219) were shared between *S. nigrum* and *S. dulcamara* only (Table S5 Fig. 3), while 27 orthogroups were only present in *S. dulcamara* (0.3% or 90 genes), and *S. nigrum* had 61 unique orthogroups (1.2% or 471 genes; Table S6). The unrooted tree produced by orthoFinder (Fig. 3) grouped the *S. americanum* samples closely together and placed all Potato clade commercial crops (*S. tuberosum, S. lycopersicum* and *S. melongena*) into a clade nested within a larger clade containing *S. nigrum and S. dulcamara,* confirming previous classifications (Huang et al. 2023) and the accuracy of both genome assemblies. Susceptible and resistant/tolerant plant species/cultivars were dispersed across the tree and did not group together.

**Fig. 3.**
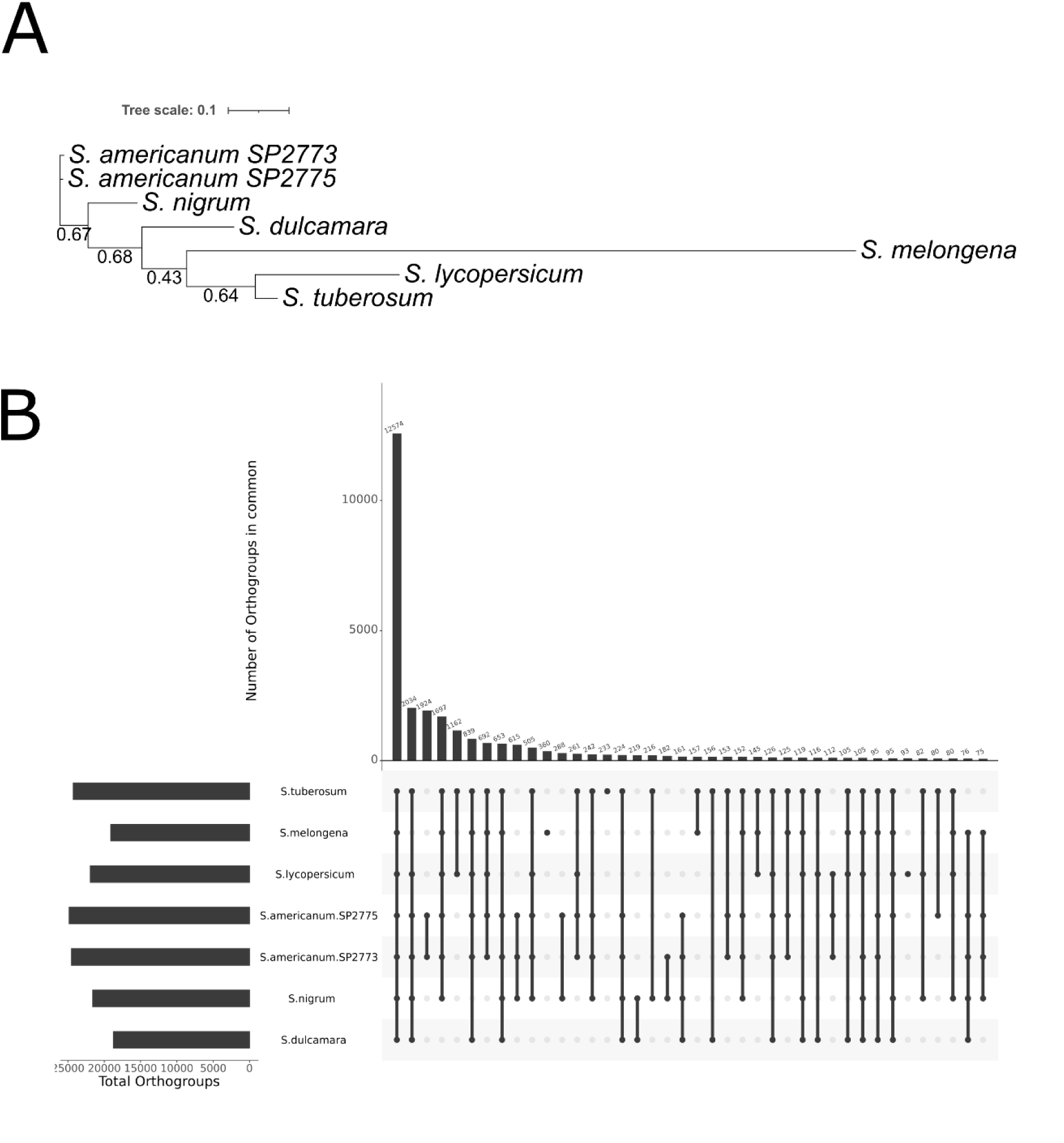
Species tree and orthogroups shared between *S. dulcamara* and *S. nigrum* and other susceptible and resistant species to *R. solanacearum*. A. Unrooted species tree was produced using the STAG (Species Tree inference from All Genes) algorithm between plant species and accessions with different susceptibility to *R. solanacearum* (resistant/tolerant species: *S. dulcamara* and *S. americanum* SP2773 and susceptible species: *S. nigrum*, *S. americanum* SP2775, *S. lycopersicum*, *S. melongena* and *S. tuberosum*) created by Orthofinder and assessing the CDS models of each specie. **B**. Orthogroups intersect between the plant species and accessiones used in A assessed with Orthofinder. Only groupings with 75 orthogroups are represented in the plot.

To identify potential mechanisms underpinning the ability of *S. dulcamara* to resist *R. solanacearum*, we first explored 90 genes belonging to 27 orthogroups that were only present in *S. dulcamara* (Table S7) using singular enrichment analysis of gene ontology (GO) terms. This revealed an enrichment of auxin transport-related GOs (Table S8). In other bacterial plant disease systems, salicylic acid defence responses stabilise auxin repressors, translating into boosted disease resistance when auxin responses are blocked (Wang et al. 2007), and during *R. solanacearum* infections, resistant tomato plants repress the transport and signalling of auxins (French et al., 2018a). Auxin accumulation, depending on PINs transport (Grieneisen et al. 2007), also explains *R. solanacearum*-related alteration of root development in the susceptible plant *Arabidopsis thaliana*, which facilitates the colonisation of the plant (Zhang et al., 2023a). The enrichment of genes associated with auxin transport in the partially resistant *S. dulcamara* may have a potential role in facilitating pathogen colonisation and establishing latent infections (Sebastià et al. 2021). However, it is also possible that these genes are involved in boosting root development and could be associated with resistance, as increases in root growth and area have been seen in resistant crops when inoculated with *R. solanacearum* (Meline et al. 2022). However, this hypothesis should be further explored with this wild plant in the future.

We also assessed the 30 genes found in common between the two resistant accessions: *S. dulcamara* and *S. americanum* SP2773 (Table S9). Here we observed: Sd_g8870, a LRR orthologous to *Arabidopsis thaliana* At1g35710; and Sd_g28574, a LRR receptor-like serine threonine-protein kinase orthologous to At1g07650, which has previously been found to be differentially expressed in susceptible potatoes when inoculated with *Dickeya solani*, the causal agent of blackleg and soft rot in potato (Hadizadeh et al. 2022). These genes are often categorised as pattern recognition receptors (PRRs) which recognize pathogen-associated molecular patterns (PAMPs) inducing PAMP-triggered immunity (PTI) in infected plants (Yuan et al. 2021). These genes could be used as targets in future works to understand their functional role during bacterial wilt disease development and confirm if they are associated with resistance/tolerance.

Overall, the comparative genomic analysis revealed a specialised group of PRRs in species that are resistant/tolerant to *R. solanacearum* and not present in other susceptible crops and wild plants. These PRRs could enable these plants to recognise and defend against a wider range of pathogens. We also identified increased numbers of auxin-signalling related genes *in S. dulcamara*, which may be linked to their role as a reservoir host to this pathogen.

### Genes common to resistant/tolerant plant species show different patterns of methylation frequency and transposable elements

After identifying PRRs only present in *S. dulcamara* and in the resistant *S. americanum* SP2773, we assessed if these potential resistance-associated genes showed higher numbers of TEs or different DNA methylation frequencies across the three genomic locales (upstream, gene body and downstream regions). As we did not have the methylation profile of *S. americanum* SP2773, we only assessed methylation frequencies and TE content in *S. dulcamara*.

We observed that Sd_g8870 and Sd_g28574 showed higher methylation in the upstream and downstream regions than other LRRs (Table S10). Sd_g8870 showed 47.6% methylation frequency in the upstream region (compared with 39.4±29.4 in other LRRs; p=0.013), whilst Sd_g28574 showed 56.9%, which is significantly higher than the average seen in other LRRs (p-value=9.6E-07). Similarly, in the downstream region of the gene, Sd_g8870 showed higher methylation than the average in other LRRs (the average in other LRRs is 47.8±28.9; p-value= 1.36E-10). Instead, the methylation frequency in the gene body was lower (27.3% for Sd_g28574, p-value=2E-10; 38.1% for Sd_g8870, p-value=0.002 in both cases than the average (48.9±29.3).

To confirm if these methylation frequencies were associated with transcriptional changes, we evaluated the expression of these specific genes and compared them with the expression of other LRRs. We hypothesized that higher methylation frequency in the promoter region would be associated with lower expression, and lower methylation frequency in these upstream regions would be associated with higher expression. In this case, the expression of these two genes was zero in roots (Fig S2B), which could be due to the higher methylation observed in upstream regions. We also observed that Sd_g8870 had a higher number of TEs than average in the upstream region (genome average is 0.24±0.60, z=,208.56, *p*<0.001; Table S10), which could be affecting methylation frequency and consequently the gene expression. However, as we measured the methylation frequency using leaves, future experiments should look to see if the methylation profiles of other tissues, such as root, correlate with the methylation frequency measured in leaves.

Overall, these results could indicate the expression of redundant disease-related genes, especially in roots, being the point of entry of *R. solanacearum,* is being suppressed by DNA methylation in upstream regions. These results are in line with the results seen by Kong et al. (2018) who suggested other disease resistance (*R*) genes such as nucleotide-binding leucine-rich repeat expression in *Arabidopsis* are regulated by DNA methylation, predominantly found in the promoter and gene body. It is also likely that these genes are tightly epigenetically controlled until needed, maintaining high methylation frequency until the plants are infected by *R. solanacearum*. This would be similar to the reduction in methylation of defense-related genes to enhance resistance in *Arabidopsis* when it responds to *Pseudomonas syringae*’s PAMPs (Lee et al. 2023), Unfortunately, we cannot judge whether this is the case using our current data, as it was generated from uninfected plants, so future work should focus on comparing expression and methylation profiles of these genes before and after inoculation with *R. solanacearum*.

Within the genes in common between *S. dulcamara* and the resistant *S. americanum* SP2773, we also found the uncharacterised Sd_g2132, which had 3 and 4 LTRs in the upstream and downstream regions respectively (compared to genome averages of 0.24±0.60, z=758.85 *p*<0.001; and 0.22±0.56, z=1115.67 *p*<0.001, respectively). We also saw a higher methylation frequency of this gene in all genomic locales than the average considering all genes in the genome (Table S10; the average was 24.8±32.8%, 26.9±35.11%, 24.9±32.9% for upstream, gene bodies and downstream regions respectively).

It is possible that the TEs associated with the upstream region of Sd_2132 could have a role in the transcription of this gene and to identify the role of this gene, we checked its orthologues and found it belongs to the orthogroup OG0024616, which also contains two *S. americanum* genes (sp2273chr03_g00625.1 and sp2273chr08_g01052.1) and the *S. dulcamara* gene Sd_g5379.2. was identified as SWI SNF-related matrix-associated actin-dependent regulator of chromatin subfamily A containing DEAD H box. Sd_g5379.2. also showed 89.14% identity (E-value 7E-69) to the *Capsicum annum* target of rapamycin complex (TOR) subunit LST8-1 (LOC107863575) (GenBank ID= XM_047408420.1). The TOR pathway is a central regulatory network conserved across kingdoms (Ryabova et al. 2019). In plants, it is involved in regulating gene expression and other metabolic adjustments controlling the switch between stress and growth and has been associated with responses to nutrients, and stresses such as pathogens and hormones (Ryabova et al. 2019).

Interestingly, when the *R. solanacearum* RipA5 (former AWR5) effector was expressed in yeast, this caused effects resembling rapamycin’s inhibitory effect on TOR signalling and *in planta* this effector caused a decrease in the activity of enzymes regulated by TOR (Popa et al. 2016). Popa et al. (2016) concluded that TOR-deficient plants are more susceptible to *R. solanacearum,* and that the bacterium inhibits this pathway. Our results suggest Sd_g2132 could be important during bacterial wilt infections and its transcription regulation could be associated with higher methylation and higher TE content.

Altogether, these analyses identified potential epigenetic associations in genes found in common in plant species resistant to *R. solanacearum*.

## Conclusions

We have assembled and annotated two new wild plant species genomes: *S. dulcamara* and *S. nigrum* which showed different susceptibility to the major plant pathogen *R. solanacearum*. We have assessed the methylation frequency and role of TEs across the whole genome in both species and confirmed *S. nigrum* had a higher methylation frequency than *S. dulcamara*, which is probably associated with a higher TE content and methylation. We identified auxin-transport-related genes that are only present in *S. dulcamara*, which we hypothesise could be associated with its role as a reservoir plant for *R. solanacearum* or to resistance mechanisms. PRRs identified in common between the resistant/tolerant S*. dulcamara* and *S. nigrum* SP773, and not present in susceptible crops or wild species, could also be a target for future work to validate their functional roles during disease development. Higher methylation frequency of these PRRs was also associated with lower gene expression in some plant tissues, suggesting epigenetic regulation of genes involved in disease resistance. The improved genome resources and candidate genes provided here will facilitate further investigations into the defence against this important bacterial pathogen.

## AUTHOR CONTRIBUTIONS

SFO extracted the DNA and RNA, assembled and annotated both genomes, performed the comparative genomics and wrote the first version; SJ and LG assessed the quality of the DNA and RNA and ran the PromethION; KH performed the flow cytometry analysis, JD and HS provided expert horticultural support; VPF obtained funding, ALH obtained funding, coordinate the project, review and edited the manuscript. All authors have read and approved the manuscript.

## Data availability statement

The genomes are available under the NCBI BioProject: PRJNA1014230 and PRJNA1017508, including raw data, and genome assemblies (JBBJLP000000000 and JAZKKB000000000). Genomes with full annotations and methylation calling are also available here: www/webfiles.york.ac.uk/Harper/Solanum_dulcamara and www/webfiles.york.ac.uk/Harper/Solanum_nigrum. Methylation profiles are also available in Gene Expression Omnibus (GEO) accession GSE262401 and GSE255584. Gene expression of the roots, stems, leaves, flowers and fruits is available in the GEO accession GSE283679. All the scripts used in this study can be found at https://github.com/sfortega/Solanaceae_genomes

## Acknowledgments

This project was undertaken on the University of York High-Performance Computing service: Viking and Viking2. The authors would like to thank the Bioscience Technology Facility at the University of York for assistance. We thank The Royal Botanic Gardens Kew for supplying the seeds of both nightshade species.

## Supplementary figures

**Fig. S1.**
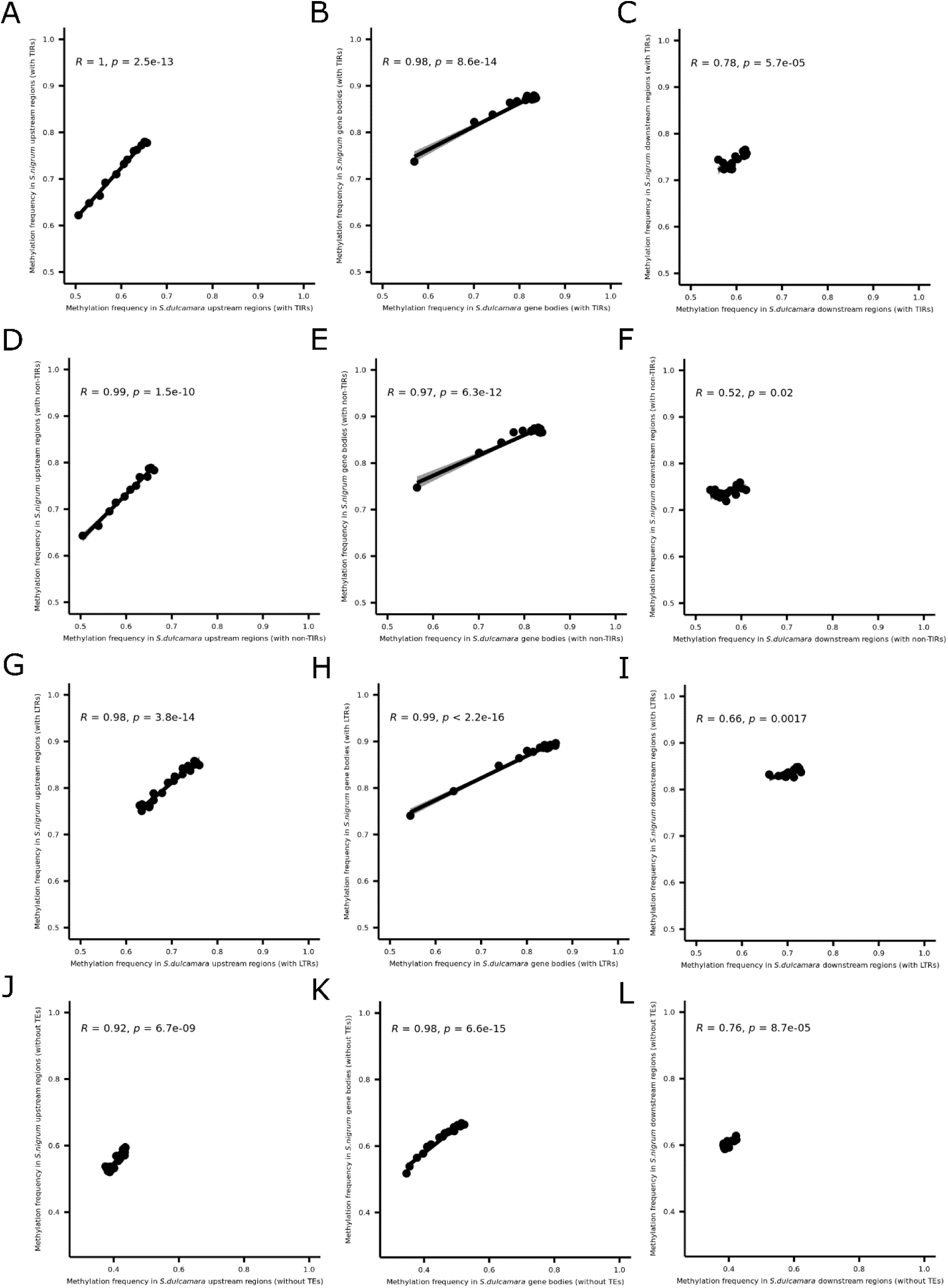
Pearson correlation between TE content in different genomic locales of the *S. dulcamara* and *S. nigrum* genes. A – C. Pearson correlation and p-value of upstream (1 kbp; A), gene body (B) and downstream (1 kbp; C) regions containing TIRs. D – F. Pearson correlation and p-value of upstream (1 kbp; D), gene body (E) and downstream (1 kbp; F) regions containing non-TIRs. G – I. Pearson correlation and p-value of upstream (1 kbp; G), gene body (H) and downstream (1 kbp; I) regions containing LTRs. J – K. Pearson correlation and p-value of upstream (1 kbp; J), gene body (K) and downstream (1 kbp; L) regions in genes without TEs.

**Fig. S2.**
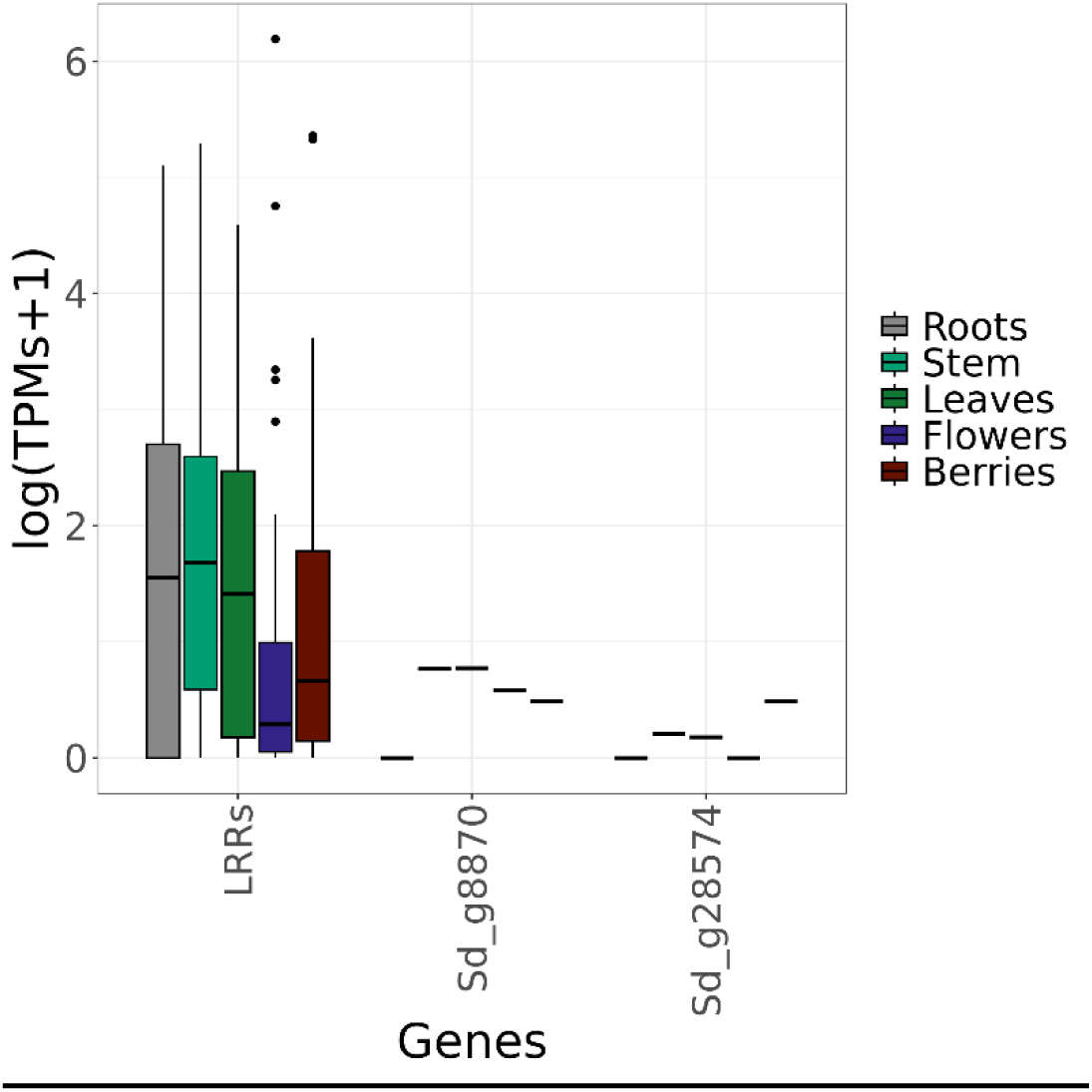
Gene expression (log (TPMs+1)) of Sd_g8870 and Sd_g28574. The expression as measured in roots (grey), stem (green), leaves (green), flowers (purple) and berries (red) of selected PRRs compared with the rest of leucine-rich repeat receptor-like kinase (LRRs).

## Supplementary tables

**Table S1**. Methylation frequencies in each contig of the *S. dulcamara* genome.

**Table S2**. Methylation frequencies in each contig of the *S. nigrum* genome.

**Table S3**. Orthogroups are assigned using SYNIMA tool. Gene names correspond with the different annotations of the genomes: “C88” corresponds with the potato genome (*S. tuberosum* C88; http://spuddb.uga.edu/data/), “Solyc” corresponds with the tomato genome (pangenome of *S. lycopersicum*; Zhou et al., 2022), “SMEL4.1” corresponds with the aubergine genome (*S. melongena*; https://solgenomics.net/ftp/genomes/Solanum_melongena_V4.1), “sp2273” corresponds wiht the resistant *S. americanum* SP2773 accession and “SaSP2275” with the susceptible *S. americanum* accession (Moon et al., 2021), “Sd” corresponds with the *S. dulcamara* annotations and “Sn” with the *S. nigrum* genome.

**Table S4**. OrthoFinder statistics of the orthogroups of the analysis with 7 plant species (resistant/tolerant species: *S. dulcamara* and *S. americanum* SP2773 and susceptible species: *S. nigrum*, *S. americanum* SP2775, *S. lycopersicum*, *S. melongena* and *S. tuberosum*).

**Table S5**. Number of common orthogroups between with 7 plant species (resistant/tolerant species: *S. dulcamara* and *S. americanum* SP2773 and susceptible species: *S. nigrum*, *S. americanum* SP2775, *S. lycopersicum*, *S. melongena* and *S. tuberosum*).

**Table S6**. Orthofinder statistics of the orthogroups and genes in each specie (resistant/tolerant species: *S. dulcamara* and *S. americanum* SP2773 and susceptible species: *S. nigrum*, *S. americanum* SP2775, *S. lycopersicum*, *S. melongena* and *S. tuberosum*).

**Table S7**. Genes in 27 orthogroups only represented by genes from *S. dulcamara*.

**Table S8**. Gene ontology enrichment of 90 genes belonging to 27 orthogroups represented only by *S. dulcamara* genes, and the genes in orthogroups in common between *S. dulcamara* and *S. americanum* SP2773, both resistant to *R. solanacearum*.

**Table S9**. Genes in 19 orthogroups in common between *S. dulcamara* and *S. americanum* SP2773.

**Table S10**. Methylation frequency across gene body, 1Kbp upstream and 1Kbp downstream for the genes in common between resistant plant species to *R. solanacearum: S. dulcamara and S. americanum SP2773*. In red, significantly higher methylation frequency than the average methylation frequency in other leucine-repeat proteins (LRRs). In blue, significantly lower methylation frequency than the average methylation frequency in other LRRs.

